# A contribution to the variability of *Prunus spinosa* L. in the vicinity of the mediaeval Castle Kolno, near Brzeg, S-W Poland

**DOI:** 10.1101/2022.12.25.521889

**Authors:** Romuald Kosina

## Abstract

In the present work, the variability of stone characteristics of mediaeval fossil forms and modern ones from a small blackthorn population in the vicinity of Castle Kolno was analysed. A modern hybrid form obtained from crossing with cultivated plums and appropriate fossil morphotypes were discovered. The analysis of the correlation of features indicated developmental relationships between them. The pattern of variability of the examined stones in the ordination space confirmed the development of the *Prunus spinosa-Prunus domestica* complex as an element of the dynamic syngameon. The population also showed variations in leaf morphotypes, which may be used in taxonomic analysis.

## 1. Introduction

*Prunus spinosa* shrubs together with other shrub species form the ecotone zone of the *Rhamno-Prunetea* syntaxon at the junction of open spaces (meadows and arable fields) and closed forest areas (Matuszkiewicz 1981; Popescu & Caudullo 2016). In the vicinity of the medieval Castle Kolno, the existing components of this syntaxon are probably remnants of an earlier phase of vegetation on the outskirts of the fields and ruins of the castle. Earlier archaeobotanical research at the site showed the possibility of crossing *P. spinosa* with *Prunus domestica* and formation of a syngameon (Kosina & Marek 2021). A hybrid taxon derived from *P. spinosa* (*Prunus fruticans* Weihe) was described from Moravia, and the key identification features were provided (Kühn 1988). Hanelt (1997) describes ×*P fruticans* Weihe as a pentaploid form derived from crossing the above-mentioned species and lists eight subspecies within *P. spinosa*.He also indicates the difficulty in differentiating the hybrid form from *P. spinosa*. The eight subspecies of *P. spinosa* highlight this challenge. Fossil specimens of plum stones, including blackthorn, are abundant, which enables to perform adequate morphometric analyses on such samples. Many medieval specimens from Douai, northern France, dated 8th to 11th c. A.D. represent the typical form of stones for the species, but also a different variety macrocarpa. It is assumed that this variety is hybrid in origin (van Zeist et al. 1994). Detailed morphometric analyses of blackthorn and cultivated plums from Denmark revealed that hybrids of both occur (Nielsen & Olrik 2001). The analysis of the variability of many traits, including stone traits, in contemporary sloe populations in the area around Wrocław provided evidence for the frequent occurrence of hybrids, particularly in populations adjacent to domestic plum crops (Staszak 2004). The present study investigated the nature of blackthorn variability in a small population in the vicinity of the medieval Castle Kolno in relation to fossil and contemporary material.

## 2. Materials and methods

The plant material analysed in this study was collected during an archaeological excavation in the former moat as well as from three jugs. Most of the fossil stones were obtained from layers at the depth of 165 to 190 cm. Stones of *P. spinosa* were described based on the following characters (codes used in the correlation matrix are given in brackets) (see also Fig. 5D):

- Length of a stone or nut (L)
- Width of a stone or nut (W)
- Height of ventral raphae for a stone (HVR)
- W/L ratio

The measured characters were selected from a larger set of data reported by Staszak (2004) for sloe stones. Each stone was treated as an operational taxonomic unit (OTU) in numerical analysis and marked in diagrams and photos by a letter and a number. Fossil stones were marked as follows: F1, F2, H1, H2, H3, H4, H5, H6, I1, K1, L1, L2, and M1, while modern ones followed Ps1 to Ps20. Ps1 and Ps12 to Ps20 were collected near the Budkowiczanka River, and they are of putative hybrid origin. The specimens Ps19 and Ps20 were stones from unripe and dried fruits. The other Ps specimens were collected nearby the castle hill and near the Stobrawa River. The current analysis is complementary to the previous one (Kosina & Marek 2021). The analysed collection of stones included all fossil specimens and contemporary specimens collected in 2022 (Fig. 1). Compared to the previous analysis, the current one did not include specimens collected elsewhere in Poland. Thus, the current set represents one population with narrowed variability. In this population, more vegetative reproduction (clones) than generative reproduction (small number of fruits on bushes) was observed (Fig. 2).

**Fig. 1.**
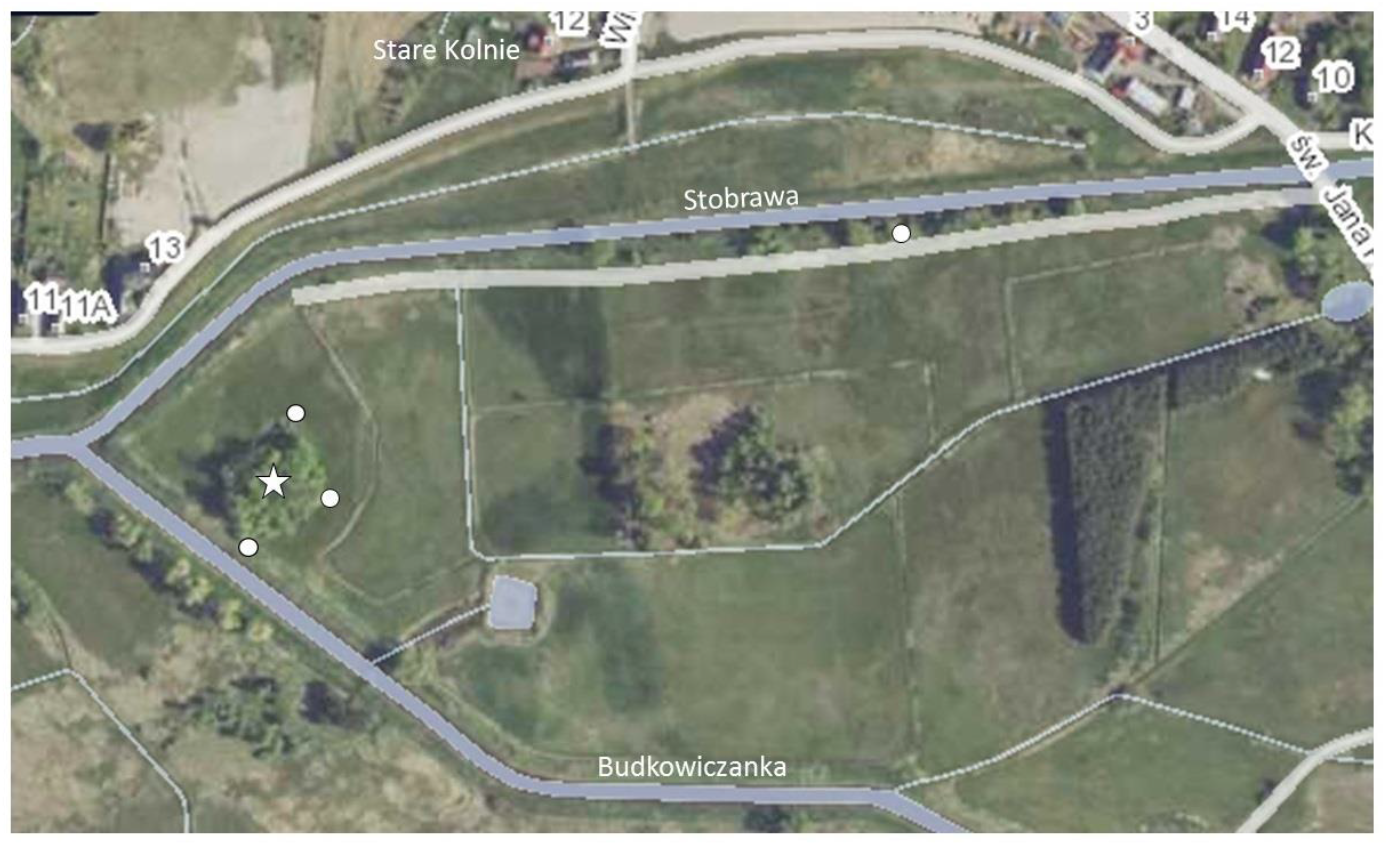
A satellite map of the Castle Kolno area. The hill of the castle covered with trees (star) and blackthorn fruit collection points (white dots) - N and E edge of the hill and over Stobrawa and Budkowiczanka rivers. The hill is located at the confluence of the rivers. http://pzgik.geoportal.gov.pl/prng/Miejscowosc/PL.PZGiK.204.PRNG.00000000-0000-0000-0000-000000129067-100262

**Fig. 2.**
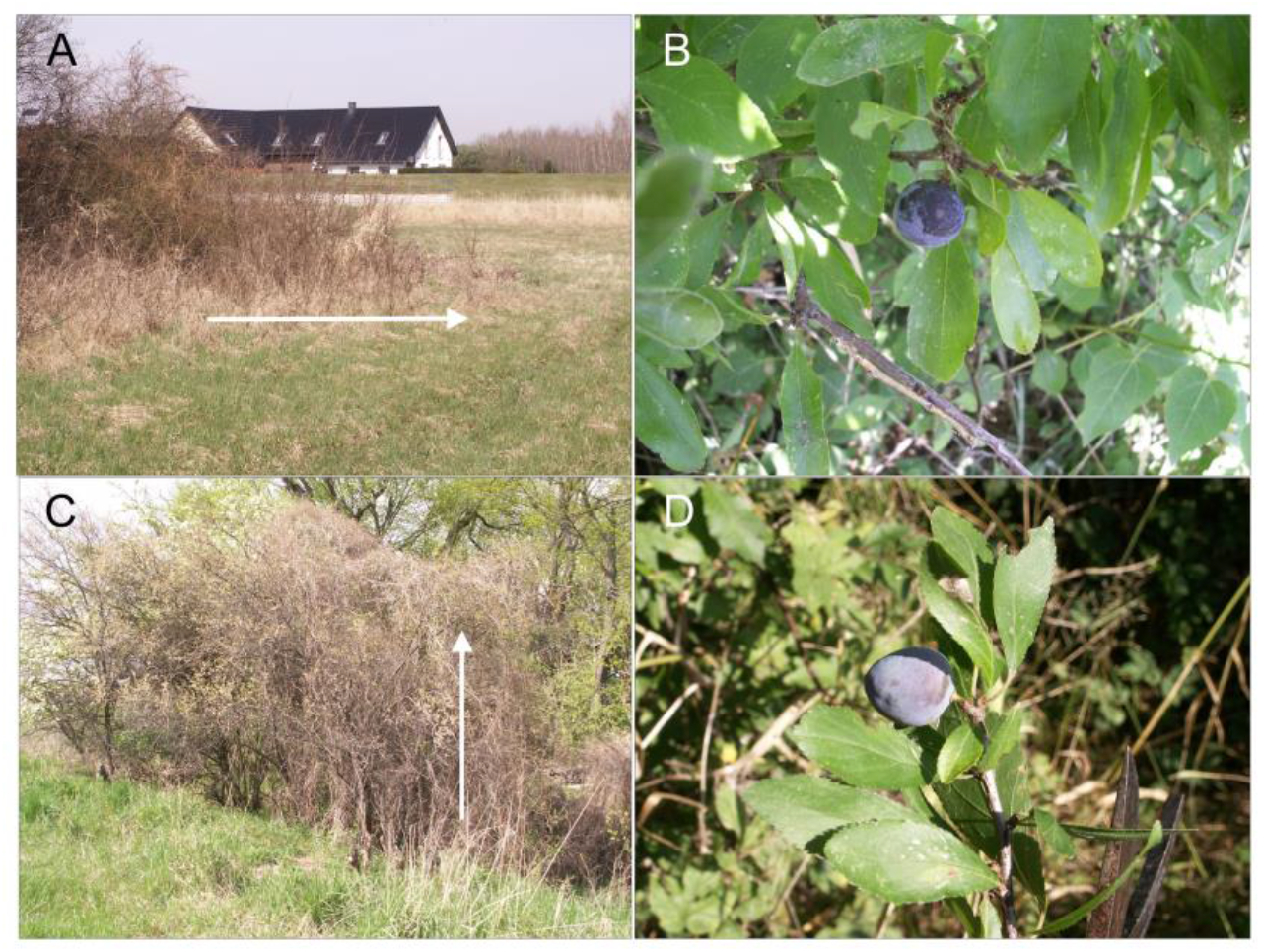
Shrubs of *Prunus spinosa* in the vicinity of Castle Kolno. A – a specimen on the N side of a hill with a clear clonal growth (arrow), B – a short stem of a specimen of a wild form near the Stobrawa River, C – a hybrid specimen near the Budkowiczanka River showing a clear vertical growth (arrow), D – a short stem of a hybrid specimen.

A matrix of average taxonomic distances between OTUs (*n* = 33) within a given set of OTUs was generated. This matrix was transformed into a configuration matrix using the Kruskal’s method of nonmetric multidimensional scaling (nmMDS), and the configuration matrix was later applied to position the OTUs in a minimum spanning tree (MST) in a three-dimensional (*x*, *y*, *z*) coordination space (see Figs. 3 and 4). Numerical analyses were performed using NTSYS software (Rohlf 1994).

**Fig. 3.**
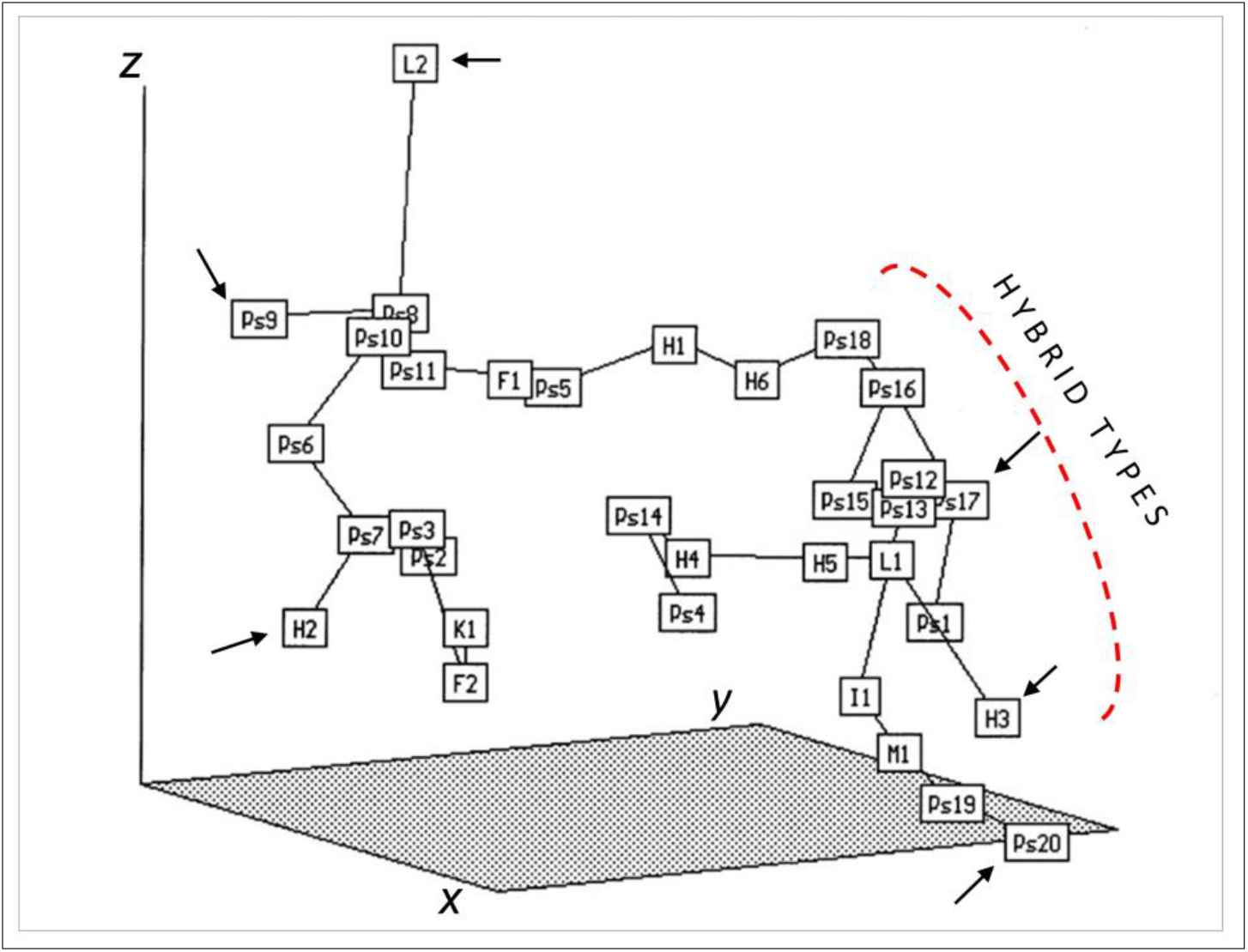
Minimum spanning tree of the contemporary and fossil stones (OTUs) of *Prunus spinosa* in an ordination space (*x*, *y*, and *z* axes). OTUs were described by four traits of the stones. Extreme OTUs are marked by arrows. Putative hybrids between *P. spinosa* and domesticated plums are marked by a broken red line. Abbreviations are provided in ‘Materials and Methods’.

**Fig. 4.**
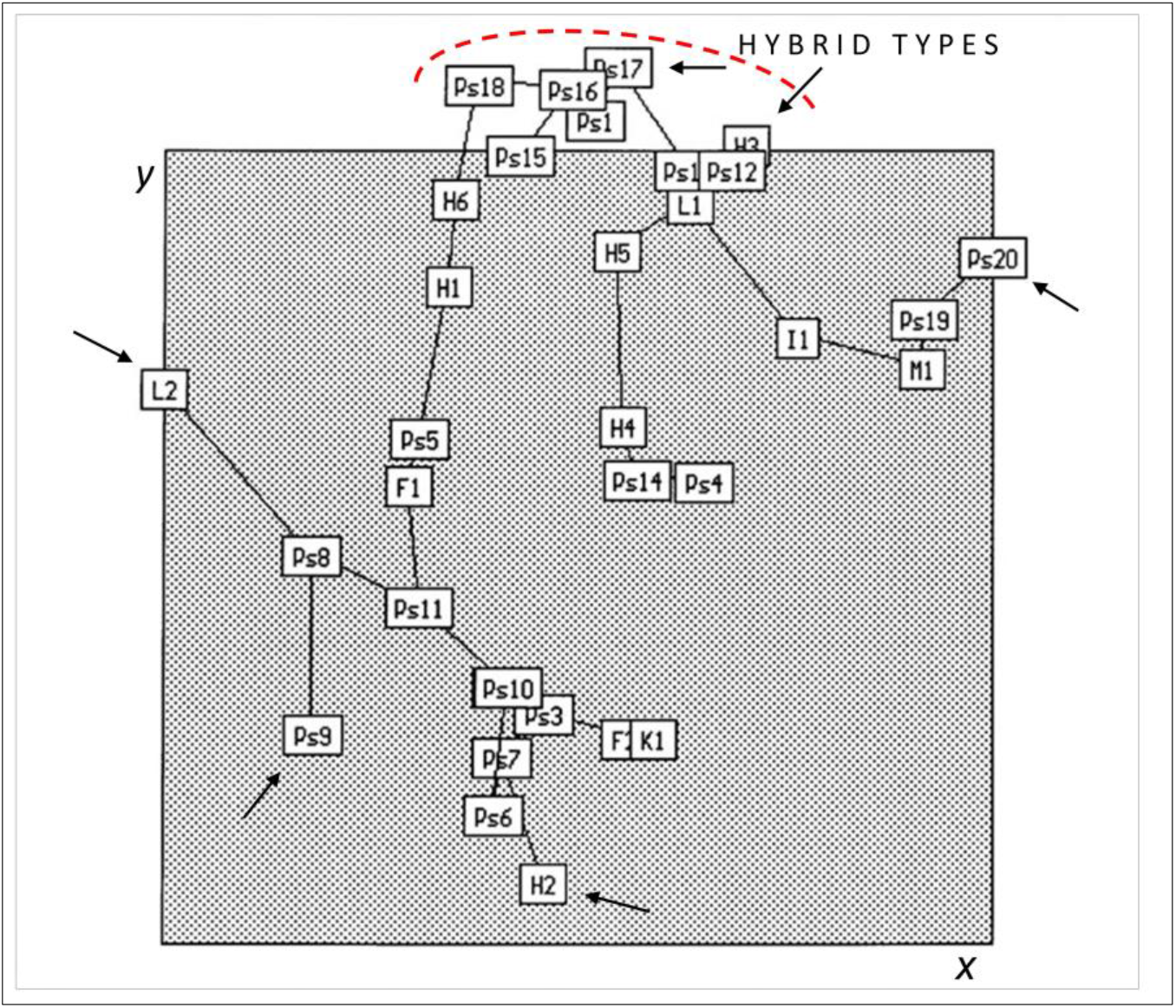
An aerial view of MST from Fig. 3. OTUs are viewed along the ordination *x* and *y* axes.

**Fig. 5.**
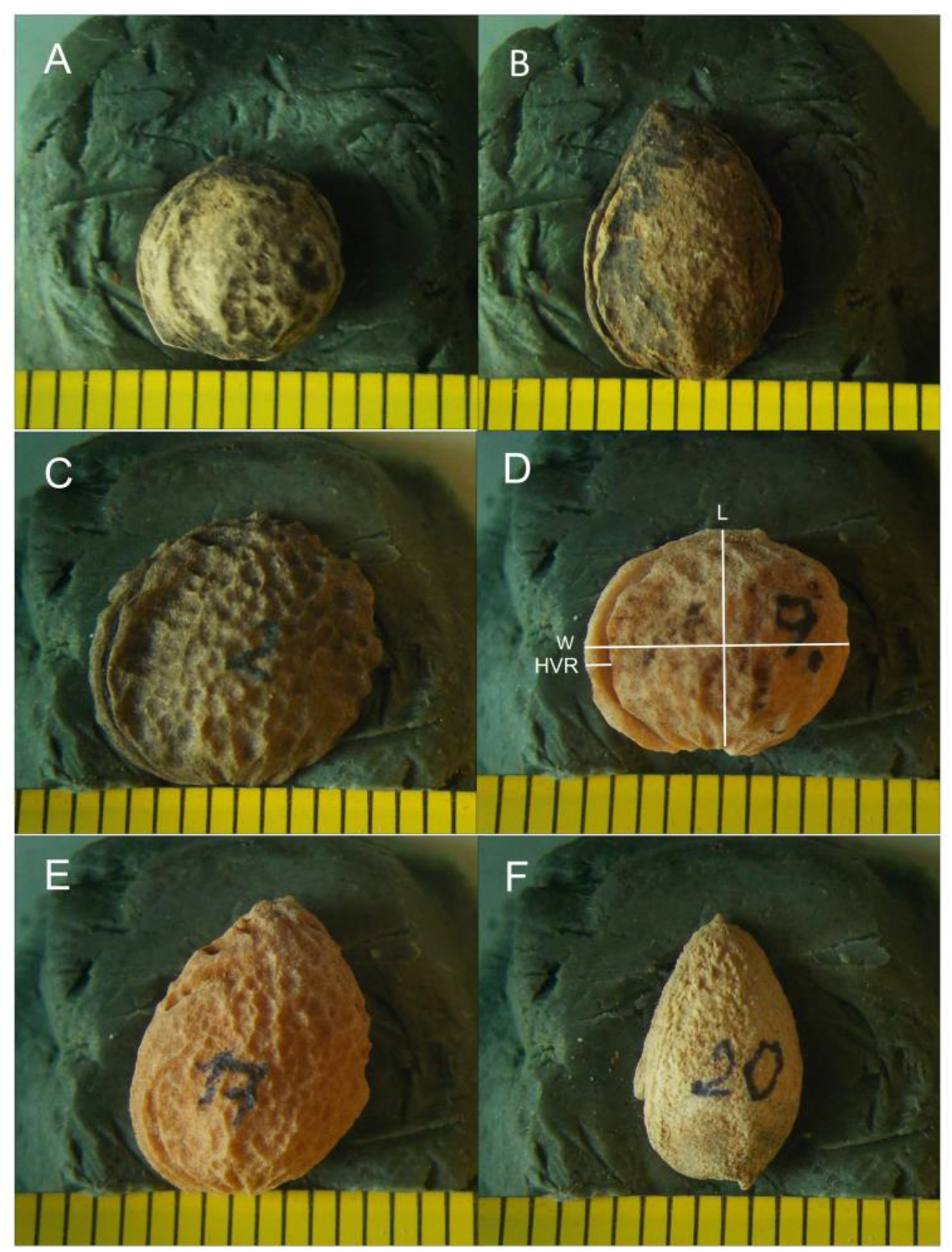
Morphotypes of fossil and contemporary stones of *Prunus spinosa* occupying extreme positions in the minimum spanning tree (see Figs. 3 and 4). A - H2; B - H3; C - L2; D - Ps9; E - Ps17; F - Ps20. Measured traits of stone are marked in D. Scale in mm.

## 3. Results and discussion

### 3.1. Correlation analysis

The analysed collection of 33 stones was derived from the local population, which included both fossil and modern ones. These stones are not from identical populations, although they may appear to be from the same origin. The previously analysed set (Kosina & Marek 2021) represented broader inter-stone variability. This situation may be characterised by several and larger significant correlation coefficients. A significant correlation was observed between the two stone dimensions, namely L and W. Presently, as shown in Table 1, this coefficient is non-significant. The remaining significant coefficients clearly indicate the relationship between the development of the stone, but do not indicate inter-population genetic variability. This conclusion confirms that the fossil and modern populations are genetically close (similar development).

**Table 1.** Pearson’s coefficients of correlation of stone characters of *Prunus spinosa* (*n* = 33)

| Characters | L | W | HVR | W/L |
| --- | --- | --- | --- | --- |
| L | 1.00 |  |  |  |
| W | 0.16 | 1.00 |  |  |
| HVR | 0.14 | 0.33* | 1.00 |  |
| W/L | -0.67*** | 0.62*** | 0.13 | 1.00 |

\*, \*\*\* - significance level at $\alpha = 0.05, 0.001$ , respectively

### 3.2. Ordination analysis

The extreme OTUs in the MST (Figs. 3 and 4) were as follows: H2 (Fig. 5A) - small, round, cherry stone-shaped stone; H3 (Fig. 5B) - large stone, tall with a sharp tip, fully developed, possible hybrid morphotype; L2 (Fig. 5C) a stone with a clear texture, almost round, similar to the modern *Prunus domestica* subsp. *syriaca*; Ps9 (Fig. 5D) - a specimen from the hill of the castle, with width greater than height; Ps17 (Fig. 5E) - a specimen from a hybrid specimen near the Budkowiczanka River, almost identical to the fossil specimen shown in Fig. 5B; types Ps19 and Ps20 (Fig. 5F) - stones from fruits that died at an early stage of development due to drought, elongated, with a sharp tip, similar to types H3 and Ps17.

The morphotype analysis of the leaves of the short shoots (Fig. 6) showed that the hybrid type had larger leaves, although it was morphologically similar to the types from the hill of the castle and from near the Stobrawa River. The leaves of this hybrid type were not similar to the leaves of *P. domestica.* Moreover, the fruits of the hybrid type were larger and had longer pedicels (Fig. 2B,C) than those of the typical *P. spinosa.*

**Fig. 6.**
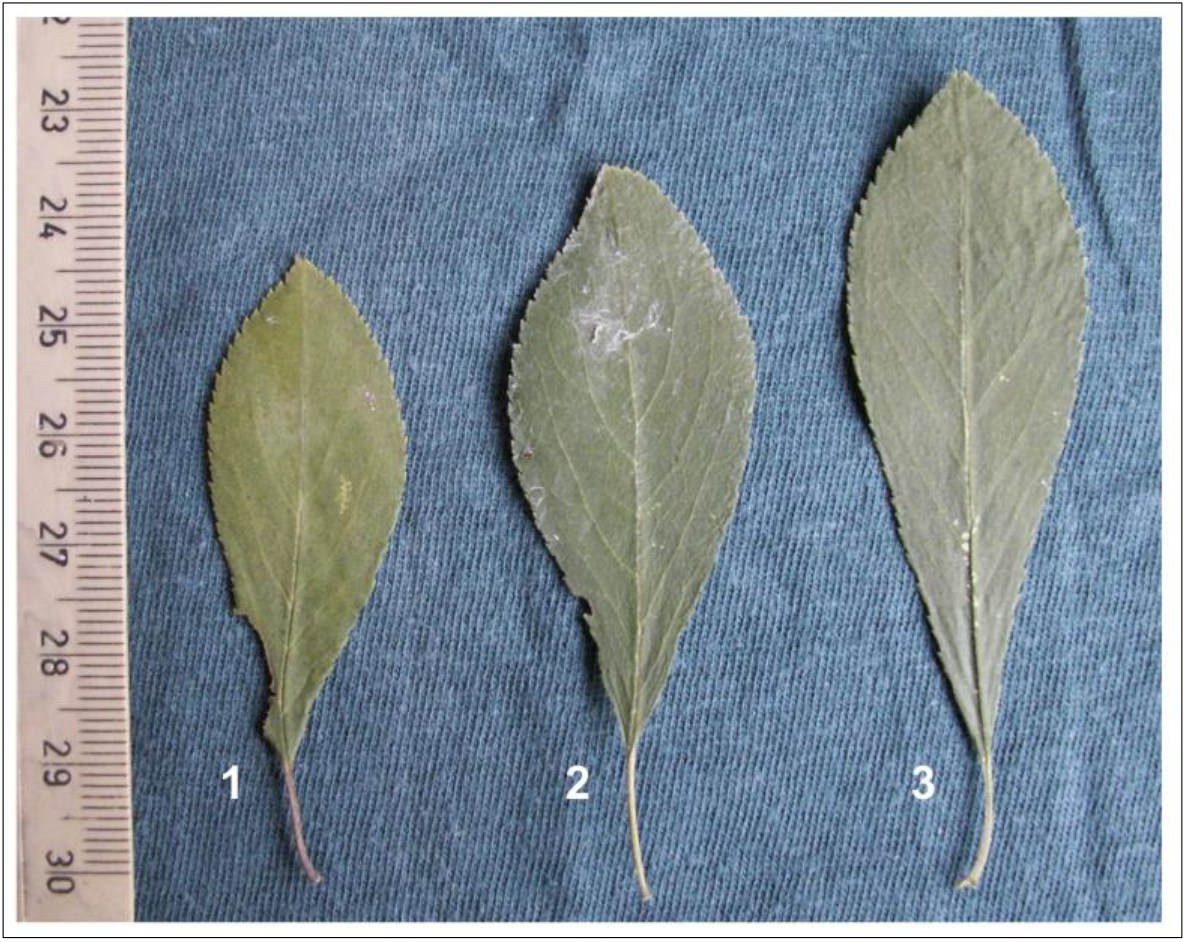
Morphotypes of leaves of wild and ‘hybrid’ forms of *P. spinosa* in the vicinity of Castle Kolno. 1 - a specimen near the Stobrawa River, 2 - a specimen from the outskirts of the castle hill, 3 - a ‘hybrid’ specimen near the Budkowiczanka River. Scale in mm (cm).

These findings indicate that in the vicinity of Castle Kolno, both in the Middle Ages (14th-15th c.) and at present, the wild blackthorn bushes also include types with the characteristics of a hybrid between sloe and cultivated plums. Hybrid shrubs-trees are characterised by erectoid growth, weak clonal reproduction, and larger and elongated fruits. A specimen of the dried ripe fruit of the hybrid is approximately 13.5 mm long with an approximately 3-mm-long pedicel. These dimensions are below those reported for ×*Prunus fruticans* by Kühn (Kühn 1988). Hanelt (1997) indicated the difficulties in differentiating hybrids in *P. spinosa* because of the high variability within the species. Fossil types reported by van Zeist et al. (1994) and described as *P. spinosa* var. *macrocarpa* vary in size and shape they are round or ovoid, but without the conical, sharp tops similar to that of *P. domestica*. A slightly greater variability of fossil blackthorn stones from a site near Heidelberg from the 13th century was described by Körber-Grohne (1979). For large mediaeval samples of *P. spinosa* from Lübeck and modern var. *macrocarpa* from Schleswig-Holstein, the variability of the stone features of the modern form showed a broad distribution, while that of the fossil type form showed a narrow one (Kroll 1980). This finding may suggest recombinational variation and hybrid origin of the modern var. *macrocarpa*.

Here, it should be emphasised that distinct hybrid types in the area around Wrocław, southwestern Poland, were found in the vicinity of former settlements or existing plum plantations (Staszak 2004). Because the spectrum of subspecies and cultivars of cultivated plums is wide and because cross-breeding with *P. spinosa* is a sporadic but constant process over time, the hybrid types show different morphotypes. Fossil type classified by Pollmann et al. (2005) as *Prunus insititia/spinosa* from the 3th c. A.D. from a Roman settlement in Switzerland was almost identical to the modern type shown in Fig. 5D, which is similar to the fossil ‘*syriaca*’ specimen shown in Fig. 5C.

The modern populations of *P. spinosa* show a large variability of traits; however, the traits of the stones were considered as stable and valuable for taxonomic analysis (Hübner & Wissemann 2004; Staszak 2004). Covariance analysis of the characteristics of the stones and leaves of this species confirmed that the characteristics of both organs can be used independently in such analyses (Kosina 2005).

A detailed morphological and phenological analysis (Vander Mijnsbrugge et al. 2016) and breeding tests (Woldring 1997/1998) of the *P. spinosa* – *P*. × *fruticans* complex showed that the latter taxon is certainly a hybrid of blackthorn and cultivated plums as a result of various crossings. The clear uniparental dominance of traits in the principal components analysis (PCA) diagram for Denmark populations of *P. spinosa* and *P. domestica* ssp. *insititia* indicated that backcrosses of putative hybrids occurred more frequently with *P. domestica* (Nielsen & Olrik 2001). According to Grant (1981), these data reinforce the view that the wild-cultivated plums complex is a dynamic syngameon in the speciation process (Kosina & Marek 2021). The stigma-pollen responses between *P. spinosa* and *P. cerasifera* (Staszak 2004) justify the inclusion in the syngameon of plums of the latter species present in Europe since the early Middle Ages.

## 4. Concluding remarks

Fossil forms from the Middle Ages and contemporary *P. spinosa* stones collected near the medieval Castle Kolno indicate the presence of hybrid types originating from spontaneous crossings with domestic plums in this area. These stones can be related to the taxonomically recognised type *P* × *fruticans*. Correlation analysis of stone features indicates that these dependencies are developmental and are already manifested in immature forms. The pattern of OTUs variation in the ordination space for a narrow local population corresponds well to that of the syngameon component. Plum orchards probably existed in the nearby settlement of Stare Kolnie in the Middle Ages, which allowed crossings between the cultivated plums and blackthorn. The determination of whether a recognised hybrid specimen near the ruins of the castle is identical to fossil stones requires molecular DNA analysis, and the effectiveness of such analyses depends on the state of preservation of DNA in fossil remains.

## Acknowledgement

I would like to thank Dr hab. Lech Marek from Institute of Archaeology, University of Wrocław, for providing fossil material for the study.

